# Effect of Maxillary Expansion and Protraction on the Oropharyngeal Airway in Individuals with Non-syndromic Cleft Palate with or without Cleft Lip

**DOI:** 10.1101/557199

**Authors:** Najla Alrejaye, Jonathan Gao, Snehlata Oberoi

## Abstract

**Introduction:** The aim of this study was to evaluate three dimensionally the effect of the combined maxillary expansion and protraction treatment on oropharyngeal airway in children with non-syndromic cleft palate with or without cleft lip (CP/L).

**Methods:** CBCT data of 18 preadolescent individuals (ages, 8.4 ± 1.7 years) with CP/L, who underwent Phase I orthodontic maxillary expansion with protraction, were compared before and after treatment. The average length of treatment was 24.1± 7.6 months. The airway volume and minimal cross-sectional area (MCA) were determined using 3DMD Vultus imaging software with cross-sectional areas calculated for each 2-mm over the entire length of the airway. A control group of 9 preadolescent individuals (ages, 8.7 ± 2.6 years) with CP/L was used for comparison.

**Results:** There was a statistically significant increase in pharyngeal airway volume after phase I orthodontic treatment in both groups, however, there was no statistically significant change in minimal cross-sectional area in neither study nor control group.

**Conclusion:** The findings showed that maxillary expansion and protraction did not have a significant effect on increasing oropharyngeal volume and MCA in patients with CP/L.

## Introduction

Cleft palate with or without cleft lip (CP/L) is the most common congenital malformation in the craniofacial region. [1,2] Children with CP/L are known to have airway complications. [3] It has been shown through three-dimensional analysis that there is a smaller oropharyngeal height and airway volume in CP/L individuals compared with non-cleft individuals. [3]

Also, individuals with CP/L have a 30% reduction in nasal airway size compared to non-cleft controls. [2] CP/L are frequently associated with nasal abnormalities such as septal deviation, nostril atresia, turbinate hypertrophy, maxillary constriction, vomerine spurs, and alar constriction [4,5,6,7,8]. These abnormalities are due to the congenital defect itself and partly due to surgeries done to repair the orofacial defect [9,10]. The nasal abnormalities in CP/L thus lead to a reduction in the dimensions of the nasal cavity and lower airway function. [6] From reduced airway function, individuals with CP/L often have airway insufficiency, velopharyngeal incompetence, snoring, hypopnea, and obstructive sleep apnea. [2]

Introduction of Cone-Beam CT (CBCT) and imaging softwares has facilitated generation of three dimensional images for reliable assessment of the cross-sectional area and airway volume. [11] In a recent study by Karia et al., three-dimensional analysis has shown a significant reduction in oropharyngeal volume in CP/L individuals versus non-cleft individuals. [2] Another study looked at the airway volume in CP/L individuals and showed that the pharyngeal airway space is significantly more reduced in the bilateral cleft palate group greater than in unilateral cleft palate and control groups. [12] It has also been shown that the minimal cross sectional area (MCA) in the pharyngeal area is most often present in the oropharynx. If the MCA is not found in the oropharynx, it is found at the junction of the oropharynx and hypopharynx. [11]

Individuals with cleft lip and palate usually develop maxillary retrusion and crossbites after cleft repair. In order to correct this developing malocclusion, maxillary expansion and protraction is commonly used during phase 1 orthodontics for such individuals in preparation for alveolar bone grafting. It is not clear if maxillary expansion results in an increase in the volume of the oropharyngeal area. Fastuca et al. found an increase in oropharyngeal airway volume in non-cleft individuals after rapid maxillary expansion (RME) leading to greater oxygen saturation in the blood. [13] However, Zhao et al. in a retrospective study found no significant change in the oropharyngeal airway volume after RME in non-cleft individuals. [14] Palomo et al. also found no significant change in the oropharyngeal airway volume after RME. Fu et al. looked at the effects of maxillary protraction from reverse headgear and found that the pharyngeal airway volume was significantly enlarged after treatment in individuals with clefts who had protraction compared to those who did not have maxillary protraction. [15]

There is limited research on airway volume in CP/L individuals using CBCT imaging. As far as the authors know, there are no studies reported in the literature evaluating the combined effect of maxillary expansion and protraction on oropharyngeal airway in individuals with CP/L. Therefore, the aim of this study was to evaluate and compare oropharyngeal airway volume and MCA in individuals with non-syndromic CP/L using CBCT before and after Phase I orthodontics with maxillary expansion and protraction.

## Method

This is a retrospective study of CBCT data of preadolescent individuals with cleft palate with or without cleft lip (n=26) who underwent Phase I orthodontics. Written informed consent was obtained for all participants of the study which was approved by CHR. We obtained ethics approval for our study from the ethics committee at UCSF, (CHR # 10-00564). The expansion and protraction group included 18 individuals (11 males and 7 females with CP/L; 3 cleft palate only, 5 bilateral cleft lip and palate and 10 unilateral cleft lip and palate), (ages, 8.4 ± 1.7 years). Individuals had initial and final CBCT scans taken with Care Stream (CS 9300, Carestream Health, Inc, Rochester, NY, USA) as part of their orthodontic treatment. The scans were stored in a DICOM (Diagnostic Imaging and Communications in Medicine) format file and loaded into the 3dMDvultus software (Atlanta, GA) for 3D airway analysis.

All individuals included in the expansion and protraction group had maxillary expansion with a fan-shaped or hyrax expander and protraction with a face mask as part of phase 1 orthodontics. Average length of treatment (observation) was 24.1± 7.6 months. The individuals with CBCT scans that were distorted or not showing the superior tip of the epiglottis clearly or other important landmarks were excluded. The control group included 8 (3 males, 5 females with CP/L; 3 cleft palate, 5 unilateral cleft lip and palate) individuals that had teeth alignment only with the inclusion criteria of non-syndromic CP/L who had no maxillary expansion, protraction, or prior orthodontic treatment.

For ethical reasons, we conducted a retrospective study utilizing the data derived from the computed tomography database of the UC San Francisco Orthodontic Clinic.

Selection of patients for the cohort selection was done by searching for patients that qualify with our inclusion criteria and utilizing all individuals that have qualifying CBCT scans. Bias in this selection process can be found in an uneven matching of gender between the treatment and control group.

After loading each CBCT scan into the 3dMDvultus software, the scan was oriented with the palatal plane parallel to the horizontal plane in the sagittal view to standardize the analysis. The oropharyngeal airway extending from the palatal plane to the superior tip of the epiglottis was outlined visually by a single investigator. (Figure 1) Cross-sectional areas were calculated for each 2-mm distance over the entire length of the airway. Measurements of the total oropharyngeal airway volume and minimal cross-sectional were calculated before and after Phase I orthodontic treatment using 3dMDvultus software (3dMD, Atlanta, GA). To determine reliability, we measured the airway volume and MCA in five random individuals after two weeks of completing the initial measurements utilizing the described measurement methods and compared the measurements to the initial findings. Changes in airway volume and MCA within each group were analyzed using matched pairs Wilcoxon signed-rank test while the changes between the two groups were compared using independent 2 sample Mann–Whitney U (Wilcoxon rank-sum) test. The data were analyzed using JMP (version 14) software (SAS Institute Inc., NC, USA) at a level of significance of 0.05.

**Figure 1.**
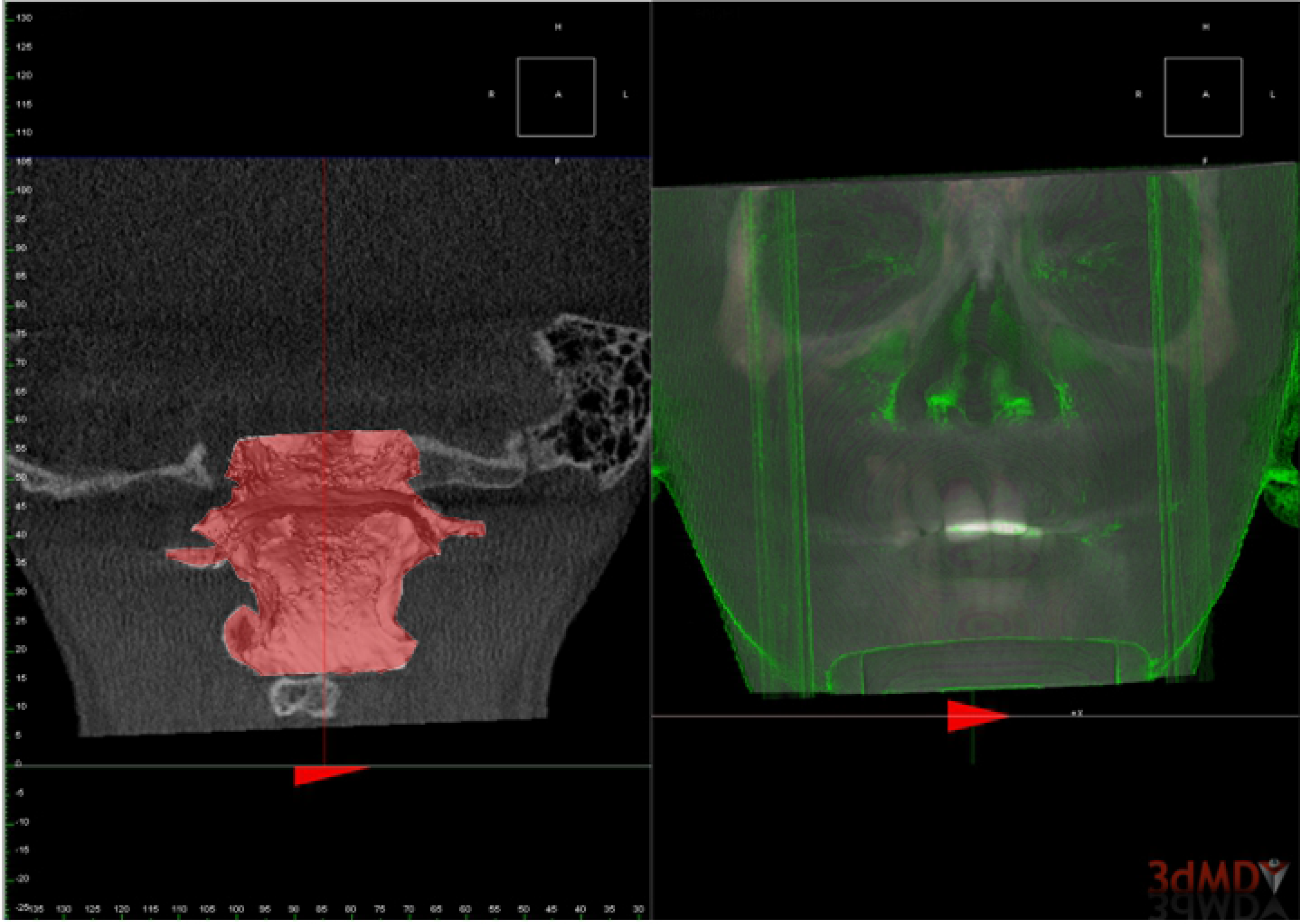

## Results

The age distribution and observation duration (24 months) between the study and the control group showed no significant differences utilizing an independent sample t-test (Table 1) Fisher exact test showed no significant difference regarding the male to female distribution amongst the two groups. (Table 2)

**Table 1:** Comparision of age and treatment duration of sample sets.

|  | Expansion + Protraction |  | Control |  |  |
| --- | --- | --- | --- | --- | --- |
|  | Mean (range) | SD | Mean (range) | SD | P-value |
| T0 (y) | 8.4 (6.8-12.8) | 1.7 | 8.9 (7.9-10.8) | 1 | 0.64 |
| T1 (y) | 10.4 (8.2-14.5) | 1.9 | 11 (8.9-12.7) | 1.7 | 0.82 |
| Duration (mo) | 24.1 (15-42) | 7.6 | 23.4 (10-50) | 13.9 | 0.4 |

**Table 2:** Gender distribution of the sample sets.

|  | Male | Female | Total | P -value |
| --- | --- | --- | --- | --- |
| Expansion/protract. | 11 | 7 | 18 | 0.4 |
| Control | 3 | 5 | 8 |  |
| Total | 14 | 12 | 26 |  |

The method of measurement of the oropharyngeal airway was found to be reliable; the intraclass correlation coefficients between the double measurements were all over 0.9.

Overall, there was a statistically significant increase in pharyngeal airway volume after phase I orthodontic treatment (P<0.05); however, there was no statistically significant change in minimal cross-sectional area within each group (P>0.05). (Table 3) The confidence interval for the change in airway volume confirmed this finding. The 95% confidence interval that the study group mean fell between [2.175,12.79]. For the control group, the 95% confidence interval was [1.21,40.83]. Since the confidence interval both do not include zero, the airway volume change in both the study and control group were statistically significant.

**Table 3:** Statistics of oropharyngeal airway volume and MCA before.

|  | Airway Volume (cm<sup>3</sup>) |  |  | MCA (mm<sup>2</sup>) |  |  |
| --- | --- | --- | --- | --- | --- | --- |
|  | T0 | T1 | Change (T1-T0) | T0 | T1 | Change (T1-T0) |
| Median | 5.8 | 8.7 | 3.6* | 58.4 | 77.7 | 20.5 |
| Median error | 0.7 | 1.2 | 0.75 | 4.3 | 6.8 | 7.7 |
| Minimum | 1.4 | 2.6 | -0.74 | 2.2 | 23.6 | -189 |
| Maximum | 49.1 | 64.5 | 34.3 | 218.1 | 356.6 | 311.3 |
| Percentile 25% (Q1) | 3.8 | 5.8 | 1.6 | 35.5 | 44.5 | -17.9 |
| Percentile 75% (Q3) | 7.3 | 15.6 | 9.8 | 96.8 | 140 | 94.7 |

The oropharyngeal airway volumes were significantly larger (p-value : <0.0001) after expansion and protraction treatment with confidence level of 95% significance at p≤.05. The oropharyngeal airway volume was also significantly larger with a confidence level of 95% (P= 0.007) for the control group with no expansion or protraction with significance at p≤.05 after treatment. (Table 4) In the expansion and protraction group, the airway volume increased 3.6 cm^3^ with a median error of 0.75 cm^3^; however, the median airway volume increase in the control group, was 2.6 cm^3^ with a median error of 3.5 cm^3^.

**Table 4:** Statistics of oropharyngeal airway volume and MCA before.

|  | Airway Volume (cm<sup>3</sup>) |  |  | MCA (mm<sup>2</sup>) |  |  |
| --- | --- | --- | --- | --- | --- | --- |
|  | T0 | T1 | Change (T1-T0) | T0 | T1 | Change (T1-T0) |
| Median | 6.9 | 10.3 | 2.6* | 84.4 | 120.4 | -16.9 |
| Median error | 1.9 | 4.7 | 3.5 | 10.5 | 7.4 | 12.6 |
| Minimum | 2.4 | 3.6 | 0.33 | 20.9 | 25.6 | -172.5 |
| Maximum | 39.8 | 76.9 | 53.7 | 216.5 | 138.3 | 101.4 |
| Percentile 25% (Q1) | 3.5 | 4.8 | 1.5 | 58.9 | 43.9 | -23.8 |
| Percentile 75% (Q3) | 8.9 | 51.6 | 38.1 | 150.2 | 128.6 | 42.5 |

The changes in volume were compared between the two groups using Mann– Whitney U (Wilcoxon rank-sum) test and no statistically significant difference detected between the two groups in either airway volume or MCA changes. (Table 5)

**Table 5:** P-Values from Change in Airway Volume.

|  | Expansion + Protraction | Control |
| --- | --- | --- |
| AWV change | <0.0001 * | 0.007* |
| MCA change | 0.12 | 1 |

Table 5: P-Values from Change in Airway Volume
|  | Expansion + Protraction | Control |
| --- | --- | --- |
| AWV change | <0.0001 * | 0.007* |
| MCA change | 0.12 | 1 |

In terms of minimal cross-sectional area, we found the 95% confidence interval to be [−9.285, 84.68] for the study group and [−95.19, 47.14] for the control group. The inclusion of zero in the confidence interval shows statistically insignificant change. The expansion and protraction group showed a median change in MCA area of 20.5 mm^2^ with a median error of 7.7mm^2^ and the control group showed a median change in MCA of −16.7 mm^2^ with a median error of 12.6 mm^2^ (Table 3). However, these changes in MCA were not statistically significantly different after treatment in either group.

## Discussion

The aim of the study was to determine the effects of maxillary expansion and protraction on the oropharyngeal airway volume and MCA in non-syndromic CP/L individuals. We measured the change in volume from initial and post treatment CBCT scans of the pharyngeal airway in children with CP/L and compared the findings to a CP/L control group who had no expansion and or protraction. To the best of our knowledge, this is the first 3-D study to measure the combined effects of maxillary expansion and protraction on airway volume and smallest cross sectional area in children with CP/L. A previous recent CBCT study assessed the change in pharyngeal airway volume in CLP individuals due to maxillary protraction without expansion. Fu et al, in that study found that the pharynx significantly increased in volume in all portions after utilizing protraction. [15] In our study we hypothesized that CP/L children with expansion and protraction had larger oropharyngeal airways post treatment compared to the control group.

According to our study, 3D imaging using CBCT and 3dMDvultus is reliable for assessing airway volume and minimal cross-sectional area.

Our data demonstrated statistically significant increase in oropharyngeal airway volume in both groups, after expansion and protraction and in the control group. Moreover, comparing the changes in airway volume between the two groups, no statistically significant difference was detected. Therefore, according to our data, there is no strong evidence that maxillary expansion and protraction itself affect orophayngeal airway volume. It could be the growth rather than expansion and protraction that caused the increase in oropharyngeal airway volume in these CP/L individuals. On the other hand, the minimal cross sectional area change was not significantly different after treatment in both expansion and protraction and control groups.

Our results were consistent with these previous studies that showed a pharyngeal airway volume increase during childhood in both non-cleft and CP/L individuals. Sheng et al reported that the pharyngeal airway depth increased from the mixed dentition stage to the permanent dentition stage in children with non-clefts over a three to four-year interval. [16] Schendel et al also reported a consistent increase in the airway volume from ages 6 to 20 years in non-cleft children. [17] Consistent with our findings, Kula et al found that children with CLP have an increase in nasal airway volume due to growth during unilateral and bilateral CP/L. [18].

However, our finding was inconsistent with Fu et al who showed no change in airway volume in CP/L individuals who did not have protraction. [15] Their study duration was about 16 months which is significantly shorter than our study and their study compared prospective experimental cleft individuals with CBCT scans to a retrospective control with Multidetector Computed Tomography (MDCT) instead of CBCT. One major difference between CBCT and MDCT devices was that individuals were examined in supine position with the MDCT and in upright position with the CBCT. These differences could have contributed to inconsistencies in the results.

With the MCA most often in the oropharyngeal area [11] and no significant change found in our study in both groups, phase one orthodontics may not have an effect on solving airway resistance problems. Growth of the airway may have an impact on airway volume size but not the MCA. [Table 3,4] The variability and no significant change in in MCA is inconsistent with findings of Abrams et al. In his study, he found that due to growth, adolescents (12-16) on average had a significantly larger MCA than children (6-11). [19] Our study is consistent

In conclusion, we found there was no statistically significant difference between the two groups in the changes in airway volume or MCA. The results of this study indicate that there is no strong evidence to show that maxillary expansion and protraction treatment have an effect on the airway volume or MCA. It is difficult or may be not possible at least for now to eliminate the effect of growth clinically to determine the expansion and protraction effect only. Future studies could provide more information by including additional assessments such as polysomnography and nasoendoscopy examinations. Also, further studies with prospective design and larger sample sizes are recommended.

## References

1. Shapira Y, Lubit E, Kuftinec MM, Borell G. The distribution of clefts of the primary and secondary palates by sex, type and location. Angle Orthod. 1999;69:523–528.

2. Canfield MA, Honein MA, Yuskiv N, Xing J, Mai CT, et al. (2006) National estimates and race/ethnic-specific variation of selected birth defects in the United States, 1999–2001. Birth Defects Res A Clin Mol Teratol 76: 747–756

3. Karia H, Shrivastav S, Karia AK. Three-dimensional evaluation of the airway spaces in patients with and without cleft lip and palate: A digital volume tomographic study. Am J Orthod Dentofacial Orthop. 2017;152(3):371–381.

4. Drettner B. The nasal airway and hearing in patients with cleft palate. Acta Otolaryngol. 1960;52:131–42.

5. Aduss H, Pruzansky S. The nasal cavity in complete unilateral cleft lip and palate. Arch Otolaryngol. 1967;85(1):53–61.

6. Warren DW, Drake AF, Davis JU. Nasal airway in breathing and speech. Cleft Palate Craniofac J. 1992;29: 511–519.

7. Hairfield WM, Warren DW, Seaton DL. Prevalence of mouthbreathing in cleft lip and palate. Cleft Palate J. 1988;25: 135–138.

8. Warren DW, Duany LF, Fischer ND. Nasal pathway resistance in normal and cleft lip and palate subjects. Cleft Palate. 1969; J 6: 134–140.

9. Gandedkar NH, Chng CK, Basheer MA, Chen PY, Yeow VKL. Comparative Evaluation of the Pharyngeal Airway Space in Unilateral and Bilateral Cleft Lip and Palate Individuals With Noncleft Individuals: A Cone Beam Computed Tomography Study. Cleft Palate Craniofac J. 2017;54(5):509–516.

10. Trindade IEK, Castilho RL, Sampaio-Teixeira ACM, et al. Effects of orthopedic rapid maxillary expansion on internal nasal dimension in children with cleft lip and palate assessed by acoustic rhinometry. J Craciofac Surg. 2010; 21:306–311.

11. Cheung T, Oberoi S. Three Dimensional Assessment of the Pharyngeal Airway in Individuals with Non-Syndromic Cleft Lip and Palate. 2012; PLoS ONE 7(8): e43405. doi:10.1371/journal.pone.0043405

12. Mordente CM, Palomo JM, Horta MC, Souki BQ, Oliveira DD, Andrade I. Upper airway assessment using four different maxillary expanders in cleft patients: A cone-beam computed tomography study. Angle Orthod. 2016;86(4):617–24.

13. Fastuca R, Perinetti G, Zecca PA, et al. Airway compartments volume and oxygen saturation changes after rapid maxillary expansion: a longitudinal correlation study [published online ahead of print February 9, 2015]. Angle Orthod. doi: http://dx.doi.org/10.2319/072014.1 ’

14. Zhao, Ying et al. Oropharyngeal airway changes after rapid palatal expansion evaluated with cone-beam computed tomography. American Journal of Orthodontics and Dentofacial Orthopedics, Volume 137, Issue 4, S71 – S78

15. Fu Z, Lin Y, Ma L, Li W. Effects of maxillary protraction therapy on the pharyngeal airway in patients with repaired unilateral cleft lip and palate: A 3-dimensional computed tomographic study. Am J Orthod Dentofacial Orthop. 2016;149(5):673–82.

16. Sheng CM, Lin LH, Su Y, Tsai HH. Developmental changes in pharyngeal airway depth and hyoid bone position from childhood to young adulthood. Angle Orthod. 2009;79(3):484–90.

17. Schendel SA, Jacobson R, Khalessi S. Airway growth and development: a computerized 3-dimensional analysis. J Oral Maxillofac Surg. 2012;70(9):2174–83.

18. Starbuck JM, Friel MT, Ghoneima A, Flores RL, Tholpady S, Kula K. Nasal airway and septal variation in unilateral and bilateral cleft lip and palate. Clin Anat. 2014;27(7):999–1008.

19. Abramson Z, Susarla S, Troulis M, Kaban L Age-related changes of the upper airway assessed by 3-dimensional computed tomography. J Craniofac Surg. 2009; 20 Suppl 1: 657–663.

20. Hakan El and Juan Martin Palomo (2014) Three-dimensional evaluation of upper airway following rapid maxillary expansion: A CBCT study. The Angle Orthodontist: March 2014, Vol. 84, No. 2, pp. 265–273.

